# Elucidating enzyme–substrate specificity through co-folding foundation model

**DOI:** 10.64898/2026.07.30.741672

**Authors:** Xiwei Cheng, Seonghwan Seo, Charlie Huh, Jihang Chen, Songlin Jiang, Pengkang Guo, Jing-Ke Weng, Woo Youn Kim, Wengong Jin

## Abstract

Enzymatic catalysis relies on precise structural and chemical complementarity, yet systematically mapping enzyme-substrate interactions remains a critical bottleneck. While structure-aware methods have advanced functional annotation, their reliance on predefined binding pockets and rigid-body docking fails to capture the ligand-induced conformational changes essential for catalytic turnover. Here we introduce Boltz2ESI, an end-to-end framework that predicts enzyme–substrate interactions by leveraging structural knowledge learned by a biomolecular foundation model. Through native co-folding, the framework inherently captures active-site plasticity without requiring predefined pocket annotations. Integrating these learned biophysical priors with global evolutionary context and geometric molecular descriptors, Boltz2ESI consistently outperforms state-of-the-art sequence-based and rigid-docking approaches. Extensive validation demonstrates that the framework accurately discriminates tight sub-family specificities, enabling effective candidate prioritization for biosynthetic pathway elucidation, as demonstrated on the withanolide pathway. Ultimately, this structure-dynamic approach establishes an actionable foundation for accelerating rational biocatalyst discovery and large-scale pathway de-orphaning.

## 1 Introduction

Enzymatic catalysis drives the chemical transformations that sustain biological systems, achieving extraordinary rate accelerations and exquisite selectivity across metabolic networks and industrial processes [1–3]. Elucidating these catalytic profiles is essential for both uncovering natural metabolic pathways and accelerating the discovery or engineering of novel biocatalysts [4, 5]. Deciphering these complex sequence–structure–function relationships remains a core challenge at the interface of chemistry, biology, and data science, where predicting the precise compatibility between an enzyme’s active site and prospective small-molecule substrates is critical.

However, experimentally mapping substrate scopes has historically relied on slow, low-throughput biochemical assays, leaving vast annotation gaps across protein databases [6]. Multi-omics approaches have partly addressed this challenge by linking gene expression patterns to metabolite profiles [7]. On the computational side, various tools have been developed, transitioning from traditional sequence alignment to sequence-based deep learning architectures [8–15]. Yet, because experimentally validated non-substrates are severely scarce, these sequence-based models generally rely on global sequence similarity [8] or evolutionary representations from pre-trained protein language models [16–18] as proxies for catalytic compatibility. By neglecting the three-dimensional spatial and chemical complementarity of the active site, such models fail to capture the functional impact of point mutations or subtle stereochemical variations that fundamentally dictate specificity. Consequently, they lack the sensitivity required to distinguish divergent substrate scopes and reactivity profiles among homologous enzymes that share the same Enzyme Commission (EC) classification [19].

The emergence of high-fidelity protein structure prediction [20, 21] has significantly enriched structure-aware approaches to protein functional annotation [22–24]. This shift has led to the development of structure-aware geometric models, most notably EZSpecificity [19] and EnzymeCAGE [4], which leverage graph neural networks and multimodal learning to map reaction coordinates to active-site structures, predicting catalytic compatibility with significantly higher fidelity than sequence-only approaches. Despite these advances, three critical hurdles limit their utility in general biocatalysis. First, these models require predefined binding pockets—typically transferred from homologous structures—which are rarely available for novel or poorly characterized enzymes. Second, they typically rely on rigid-body docking algorithms that treat the enzyme scaffold as a static entity, failing to account for the dynamic, substrate-induced conformational plasticity and induced-fit rearrangements central to the catalytic cycle [25, 26]. Finally, the scarcity of experimentally resolved enzyme–substrate complexes restricts their generalization to sequence-distant families, hindering their application to real-world tasks like de-orphaning biosynthetic pathways.

To address these limitations, we introduce Boltz2ESI, an end-to-end computational framework that harnesses the evolutionary and structural priors of a biomolecular foundation model Boltz-2 [27] to predict enzyme–substrate interactions natively through co-folding. Given only an enzyme sequence and a substrate SMILES string, Boltz2ESI operates via a two-stage structural protocol: it first localizes the active-site pocket using Multiple sequence alignment (MSA) guided co-folding, and subsequently re-folds the active site and substrate in an MSA-free regime to capture ligand-induced side-chain and backbone adaptations. By decoupling structural modeling from MSA-derived evolutionary biases, this two-step protocol forces the model to resolve the binding pose based strictly on biophysical complementarity and simulated induced-fit dynamics. From this simulated complex, the framework extracts interaction representations and a predicted structure to capture local biophysical constraints. To reconcile this local structure with global biological context, Boltz2ESI fuses these geometric descriptors with residue-level evolutionary context derived from ESM3 [28], alongside geometry-aware molecular embeddings [29] and topological fingerprints [30] that describe the substrate’s chemical environment. By integrating these complementary modalities, the framework bypasses the need for predefined pocket definitions and models the dynamic structural plasticity of the bound state directly. These multi-modal representations are subsequently blended through a geometry-conditioned interaction module featuring a four-block Pairformer stack [31] and a multilayer perceptron head. Crucially, by initializing this module with the pre-trained weights of the Boltz-2 affinity head [27], the model inherits thermodynamic priors [32].

We systematically evaluate Boltz2ESI on the comprehensive ESIBank benchmark, where it consistently outperforms state-of-the-art sequence-based and rigid-docking baselines. By learning to distinguish catalytic specificity requirements from mere ground-state thermodynamic binding affinity, the model achieves exceptional discriminative precision even among homologous enzymes sharing identical four-digit Enzyme Commission (EC) classifications. Through a detailed structural case study on citrate synthase, we demonstrate that the co-folding architecture successfully resolves large-scale conformational rearrangements—such as open-to-closed domain movements—that are physically inaccessible to static docking algorithms. Additionally, comprehensive family-level evaluation reveals that while the pretrained model generalizes robustly across diverse reaction chemistries, targeted fine-tuning provides a highly effective mechanism for adapting to previously underrepresented structural folds. Finally, we demonstrate the real-world utility of our framework by successfully prioritizing the key P450 enzymes within the complex withanolide biosynthetic pathway directly from transcriptomic candidates. Ultimately, these results establish Boltz2ESI as a powerful, generalizable tool for mapping substrate scopes, de-orphaning metabolic networks, and accelerating rational biocatalyst discovery.

## 2 Results

### 2.1 Overview of the Boltz2ESI framework

Enzyme–substrate specificity is fundamentally governed by the precise spatial and chemical orchestration within the catalytic microenvironment. To capture these delicate microenvironments, recent structure-aware computational approaches [4, 19] have advanced interaction modeling by integrating geometric descriptors of active-site pockets. However, these methods typically rely on predefined or homology-mapped binding cavities, rendering them ineffective for poorly characterized or “orphan” enzymes whose active sites remain structurally uncatalogued. Furthermore, because conventional pipelines treat the protein as a rigid scaffold and rely on post-hoc molecular docking, they fail to capture the cooperative, ligand-induced conformational plasticity that dictates catalytic selectivity across diverse enzyme families.

To overcome these barriers, we developed Boltz2ESI, an end-to-end deep learning framework that harnesses the biomolecular co-folding foundation model Boltz-2 as its structural backbone to natively capture dynamic, ligand-induced enzyme–substrate complexes (Fig. 1). Given only an enzyme sequence and a substrate SMILES string, Boltz2ESI operates via a two-stage structural protocol. The framework first localizes the putative active site using an alignment-guided co-folding simulation. Crucially, the localized pocket and the substrate are subsequently re-folded in a sequence-alignment-free regime to yield a high-resolution, unconstrained local complex. This decoupled protocol minimizes evolutionary sequence biases, forcing the model to resolve the binding pose based strictly on mutual biophysical complementarity and simulated induced-fit dynamics.

**Fig. 1.**
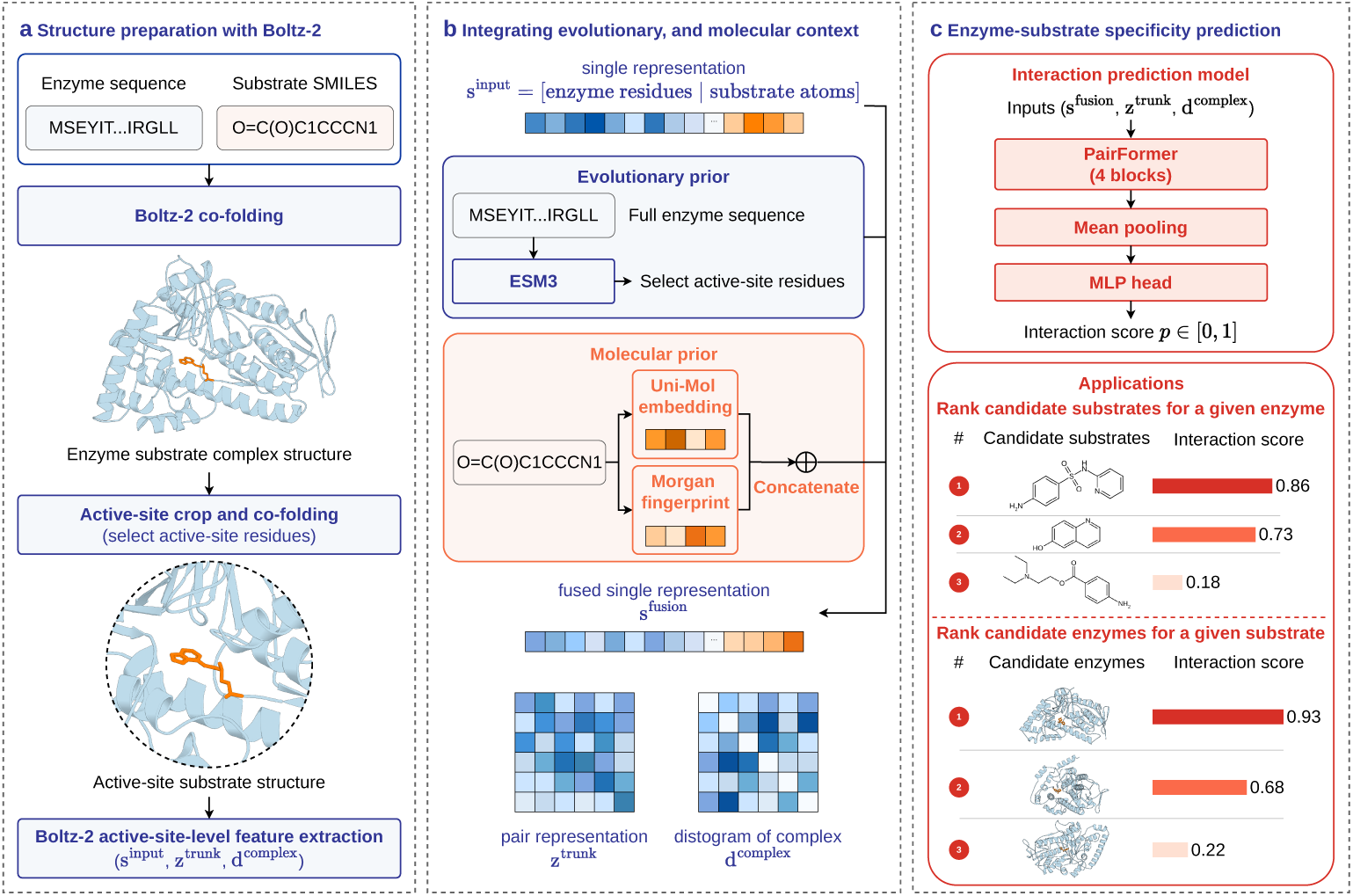
Overview of the Boltz2ESI framework. **(a)** Given an enzyme sequence and a substrate SMILES string, Boltz-2 co-folding first crops the putative active-site pocket, then the active site and substrate are re-folded without MSA, and the active-site-level single representation, pair representation, and distogram of the predicted complex structure are extracted. **(b)** ESM3 embeddings capture evolutionary context for enzyme residues, while Uni-Mol2 embeddings and Morgan fingerprints encode substrate molecular features. These priors are fused with the single representation to form a unified representation. **(c)** The interaction prediction module enriches the pair representation with the fused single representation and the distogram, processes the result through a 4-block Pairformer stack, and outputs an interaction probability via mean pooling and an MLP head. The resulting scores enable two complementary application modes: ranking candidate substrates for a given enzyme, and ranking candidate enzymes for a given substrate.

From this refined co-folded complex, Boltz2ESI extracts three complementary geometric representations that capture local physics: atomic- and residue-level identity features **s**^input^, pairwise interaction features **z**^trunk^, and spatial geometry of the predicted complex **d**^complex^. To reconcile these local structural physics with global biological context, Boltz2ESI constructs a comprehensive multi-modal feature space. It merges the structural residue features with evolutionary conservation and stability signals derived from a powerful protein language model (ESM3) [28]. Concurrently, it enriches the substrate representations by integrating geometry-aware three-dimensional molecular embeddings (Uni-Mol2) alongside topological fingerprints that describe fine-grained functional group arrangements [29, 30].

Finally, a geometry-conditioned interaction module blends these cross-modal representations through a specialized attention mechanism, restricting message passing to essential protein–ligand interfaces. By initializing this module with Boltz-2 affinity head [27], which is the pre-trained thermodynamic weights from large-scale protein-ligand complexes, the model bypasses the need to learn fundamental biophysical complementarity from scratch, significantly mitigating overfitting. The network pools these integrated features to output a final catalytic interaction score, *p*_interaction_ ∈ [0, 1], which quantitatively describes the enzyme’s capability to catalyze the target chemical transformation.

To train Boltz2ESI, we leverage the ESIBank benchmark [19], which integrates experimentally validated enzyme–substrate pairs from the BRENDA database [33] with curated family-specific datasets, totalling 323,783 high-quality pairs after quality filtering. For BRENDA-derived enzyme–substrate pairs, negative samples are constructed by pairing each positive enzyme or substrate with counterparts drawn from increasingly similar EC classes, creating a difficulty gradient from trivially distinguishable to highly challenging decoys. The benchmark is further supplemented with data from six representative enzyme families: thiolases [34], esterases [35], phosphatases [36], glycosyltransferases [37], nitrilases [38] and domains of unknown function proteins (DUFs) [39], enabling both cross-family evaluation and family-level fine-tuning (see Section 2.3).

### 2.2 Boltz2ESI enzyme-substrate specificity prediction accuracy on comprehensive test set

To systematically evaluate the capacity of Boltz2ESI to decode catalytic specificity, we conducted a comparative study using the comprehensive ESIBank benchmark. To ensure a rigorous assessment, we tested the models under two distinct validation regimes: a standard random split to establish baseline in-distribution performance, and a challenging “unknown substrate and enzyme” split designed to evaluate generalization to phylogenetically novel enzyme families and unseen chemical architectures. We benchmarked Boltz2ESI against three representative baseline frameworks: (1) ESP [40], a sequence-based architecture for generalized interaction modeling, (2) EZSpecificity [19], the current state-of-the-art structure-aware method that relies on static pocket representations and rigid molecular docking, (3) Boltz-2 [27], the base biomolecular foundation model, evaluating binding affinity predictions without downstream alignment [32]. Implementation details for baselines are provided in Supplementary Note 1. Predictive performance was robustly quantified via the area under the receiver operating characteristic curve (AUROC) and the area under the precision-recall curve (AUPR) using four-fold cross-validation.

As illustrated in Fig. 2a, where metrics are pooled over all enzyme–substrate pairs in the benchmark, Boltz2ESI consistently outperformed all baseline models across both validation settings. Within the random split, Boltz2ESI achieved an outstanding AUROC of 0.9156 and an AUPR of 0.6221, significantly surpassing the sequence-only model ESP (AUROC = 0.6572, AUPR = 0.2057) and the structure-based, rigid-docking approach EZSpecificity (AUROC = 0.8927, AUPR = 0.5995). More importantly, under the stringent “unknown substrate and enzyme” split, Boltz2ESI sustained remarkably robust predictive power, yielding an AUROC of 0.7664, outperforming both ESP (0.6778) and EZSpecificity (0.7198). Strikingly, the zero-shot affinity predictions from the Boltz-2 yielded near-random classification performance across both validation regimes (AUROC = 0.5258 and 0.5299, respectively). This performance gap highlights a fundamental biophysical distinction between thermodynamic binding affinity and catalytic specificity. While the pre-trained Boltz-2 affinity head successfully captures ground-state binding energies, it does not fully explain catalytic specificity. Consequently, non-reactive molecules that bind tightly to the pocket may be misclassified as substrates when relying solely on zero-shot affinity predictions.

**Fig. 2.**
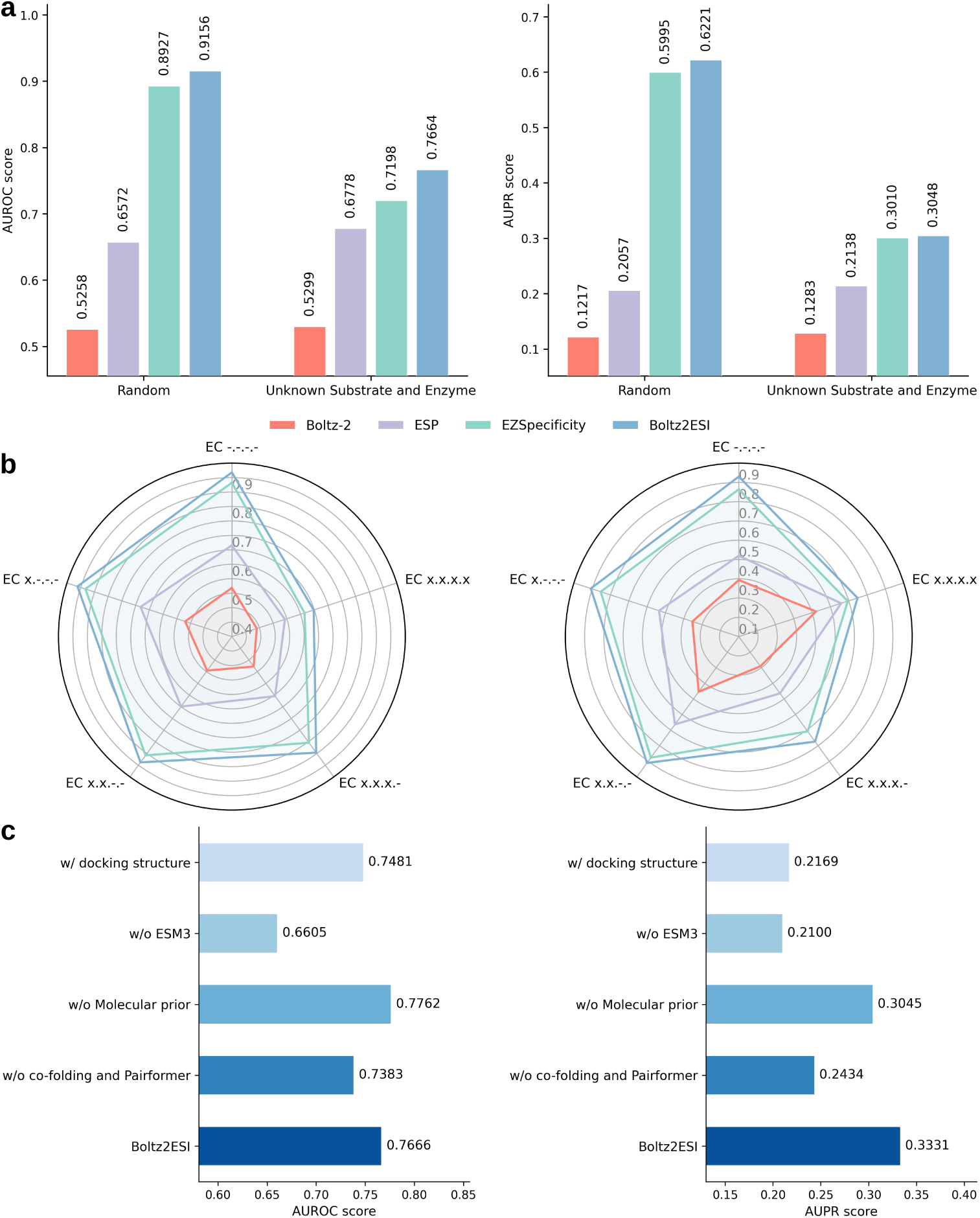
Evaluation of Boltz2ESI on the ESIBank benchmark. **a,** AUROC (left) and AUPR (right) scores of Boltz2ESI and three baselines (Boltz-2, ESP, and EZSpecificity) under the random and the unknown substrate and enzyme data splits. **b,** Discriminative resolution on the ESIBank benchmark, stratified by the number of shared EC digits between positive and negative enzyme-substrate pairs, evaluated by AUROC (left) and AUPR (right). **c,** Ablation study evaluated by AUROC (left) and AUPR (right) on the unknown enzyme and substrate split, isolating the contributions of co-folding structural representations, ESM3 evolutionary embeddings, molecular substrate priors, and co-folding structures.

By integrating co-folded structural representations with evolutionary and molecular priors and training on specificity labels, Boltz2ESI learns features associated with enzyme–substrate specificity beyond binding affinity alone, enabling more accurate discrimination of true substrates from high-affinity non-substrates.

A perennial bottleneck in biocatalysis is distinguishing true physiological substrates from highly similar molecular decoys or closely related homologous reactions. To evaluate the model’s performance at varying granularities of functional annotation, we stratified predictions across different depths of the Enzyme Commission (EC) hierarchical classification system (Fig. 2b).

As the taxonomy progresses from broad, coarse-grained catalytic classes (EC .-.-.-) down to highly specific, four-digit reaction profiles (EC x.x.x.x) and targeted families, the challenge of filtering out false positives among closely related sequences scales exponentially. Throughout this entire taxonomic gradient, Boltz2ESI consistently exhibited superior discriminative resolution compared to baseline approaches. Notably, the sustained precision at the deep EC x.x.x.x level indicates that by capturing the adaptive biophysical microenvironment within the active site, our framework effectively untangles tight sub-family specificity that remains hidden to one-dimensional sequence metrics or rigid-scaffold modeling.

To isolate the architectural components driving these performance gains, we executed a comprehensive ablation study on the first fold of the stringent “unknown substrate and enzyme” regime (Fig. 2c). Stripping the entire co-folding structural backbone and representations together with the Pairformer stack (w/o co-folding and Pairformer) decreased AUROC from 0.7666 to 0.7383 and AUPR from 0.3331 to 0.2434, confirming that geometric reasoning over the co-folded complex is integral to Boltz2ESI’s discriminative performance and cannot be compensated by sequence-level or molecular features alone. Removing ESM3 evolutionary embeddings produced the single largest performance drop across both metrics (AUROC to 0.6605, AUPR to 0.2100), demonstrating that global evolutionary context, which encodes conservation patterns, stability signals and distal allosteric effects, remains indispensable for generalizing to sequence-distant enzymes whose local pocket geometries alone are insufficient to resolve catalytic compatibility. Removal of the molecular substrate priors (Uni-Mol2 three-dimensional embeddings and Morgan fingerprints) had a more moderate effect on AUROC (0.7762) but substantially reduced AUPR (0.3045), indicating that fine-grained substrate chemistry is critical for high-precision substrate discrimination even when much of the substrate’s identity is implicitly captured within the co-folding representations. Replacing the end-to-end co-folded active sites generated by Boltz-2 with the pre-computed AlphaFold2/AlphaFill structures and AutoDock-GPU docking poses provided by EZSpecificity [19] caused a severe drop in precision, with the AUPR collapsing from 0.3331 to 0.2169. This significant degradation underscores that conventional rigid-body docking algorithms fail to capture the cooperative, ligand-induced conformational plasticity vital to real-world catalytic cycles, whereas our end-to-end co-folding pipeline naturally preserves these key structural dynamics. Collectively, these results reveal that structural co-folding and evolutionary context play complementary roles: ESM3 embeddings provide the broadest discriminative signal for distinguishing enzymes across distant families, while the co-folded complex geometry serves as the primary bottleneck for precision among closely related enzyme–substrate pairs.

### 2.3 Robust family-level performance and transferability of fine-tuning

Benchmark averages can mask uneven performance across enzyme families with disparate active-site architectures and substrate chemistries. We therefore evaluated Boltz2ESI across five enzyme families in ESIBank under the stringent unknown substrate and enzyme split, including thiolases [34], esterases [35], phosphatases [36], glycosyltransferases [37], and domain of unknown function proteins (DUFs) [39]. Nitrilases [38] were excluded owing to insufficient test data points. For each family, we report zero-shot inference with the ESIBank-pretrained model alongside a fine-tuned variant (Boltz2ESI-fine-tune) adapted from the shared checkpoint on family-specific data.

As shown in Fig. 3a, zero-shot Boltz2ESI exceeded EZSpecificity on AUROC in four of five families and on AUPR in three of five. The most consistent improvements were observed for glycosyltransferases and phosphatases, where Boltz2ESI surpassed EZSpecificity on both metrics. On esterases, Boltz2ESI and EZSpecificity achieved comparable AUROC (0.7997 and 0.8035), both substantially exceeding the sequence-based baselines, while the two methods showed comparable AUPR on thiolases (0.8216 and 0.8247) and DUFs (0.3918 and 0.4079) despite Boltz2ESI outperforming on AUROC. Fine-tuning on family-specific data yielded improvements across all families, with the most pronounced gains on thiolases (AUROC from 0.6327 to 0.7457, AUPR from 0.8216 to 0.8830) and DUFs (AUROC from 0.7372 to 0.7662, AUPR from 0.3918 to 0.5003), consistent with these families being underrepresented in the pretraining corpus rather than posing fundamental challenges for co-folding-based representations. Across all five families, fine-tuning improved AUPR in all five and AUROC in four of five (Supplementary Tables 4 and 5, Supplementary Figs. 1 and 2), confirming that ESIBank pretraining provides a broadly transferable foundation that can be readily sharpened with modest family-specific data.

**Fig. 3.**
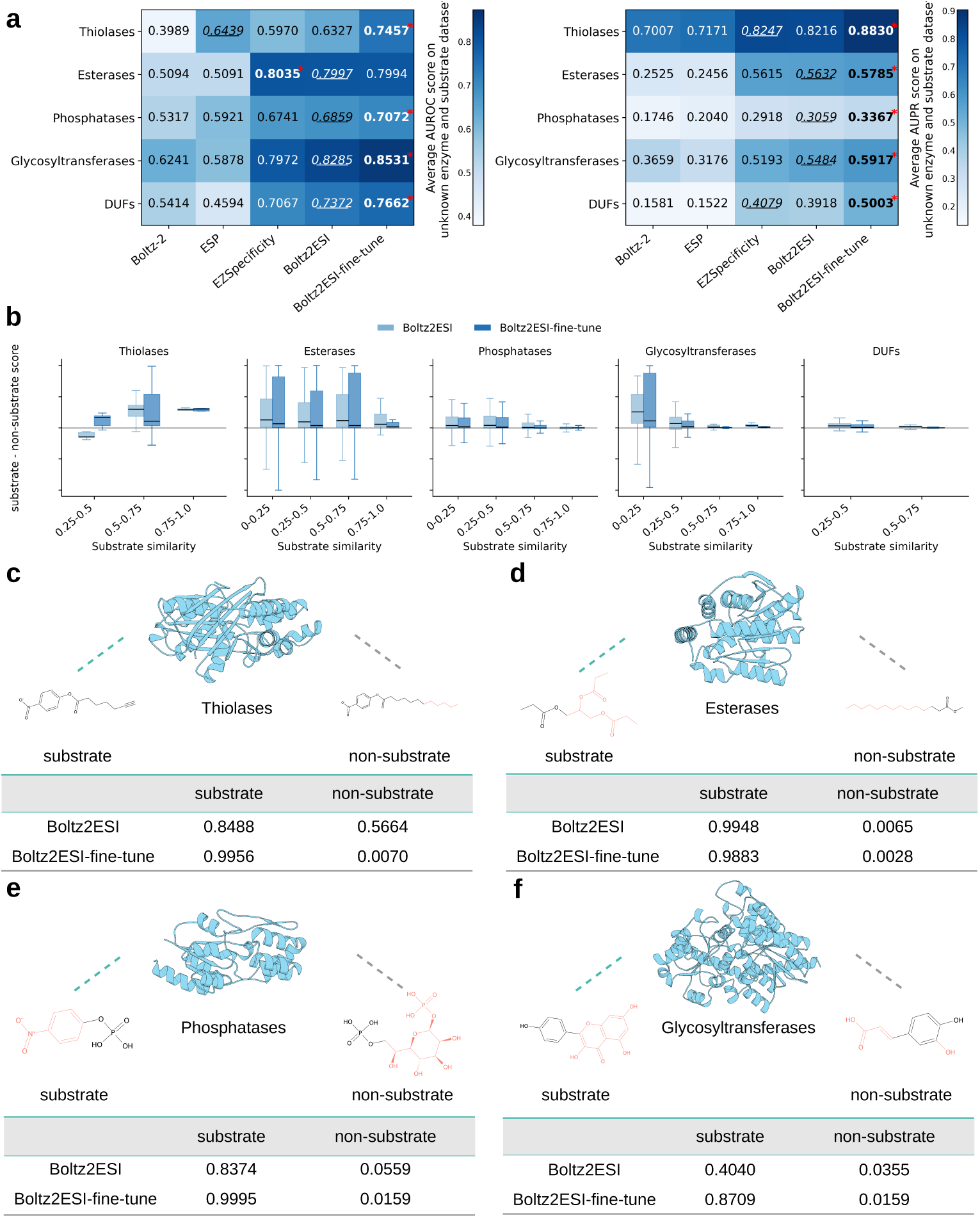
Family-level evaluation of Boltz2ESI. **a,** Per-family AUROC and AUPR heatmaps under the unknown substrate and enzyme split, comparing Boltz-2, ESP, EZSpecificity, Boltz2ESI, and Boltz2ESI-fine-tune across five enzyme families. The highest and second highest values in each row are indicated by bold and underlined text, respectively. **b,** Score margin (substrate score minus non-substrate score) stratified by Tanimoto similarity between paired substrate and non-substrate, for Boltz2ESI and Boltz2ESI-fine-tune across five enzyme families. **c–f,** Representative substrate and non-substrate pairs for thiolases (**c**), esterases (**d**), phosphatases (**e**), and glycosyltransferases (**f**), with predicted interaction scores from Boltz2ESI and Boltz2ESI-fine-tune illustrating the effect of family-specific fine-tuning on discriminative resolution. In each molecular pair, the shared scaffold is shown in black and differing substructures are highlighted in red.

To understand how fine-tuning reshapes discrimination across varying levels of substrate resemblance, we stratified the score margin (substrate score minus non-substrate score) by the Tanimoto similarity between each substrate–non-substrate pair (Fig. 3b). For esterases, phosphatases and glycosyltransferases, score margins decreased with increasing substrate similarity, consistent with the expectation that distinguishing near-identical molecular scaffolds poses a harder discrimination task. Esterases retained clearly positive margins even in the highest similarity bin (0.75–1.0), confirming that the pretrained model already captures fine-grained esterase substrate specificity. Phosphatases and glycosyltransferases showed compressed margins at high similarity but remained largely positive, indicating that pretraining provides a reasonable baseline for these families as well. Thiolases displayed the distinct pattern as score margins increased with rising substrate similarity, yet the pretrained model yielded near-zero or negative margins at low similarity levels. Fine-tuning markedly improved thiolase discrimination, with the largest gains concentrated in the low-similarity regime, possibly reflecting the underrepresentation of thiolases in the pretraining dataset. For DUFs, fine-tuning narrowed the interquartile spread of the score margin while raising the mean, a pattern consistent with the small sample size of this family, where a few outliers can dominate box-plot statistics.

Representative enzyme–substrate pairs drawn from the held-out test sets corroborate these statistical trends at the level of individual predictions (Fig. 3c–f). For thiolases (Fig. 3c), the pretrained model assigned moderate scores to both the substrate and non-substrate (0.8488 and 0.5664), yielding insufficient separation for confident classification. Fine-tuning dramatically resolved this ambiguity, elevating the substrate score to 0.9956 while suppressing the non-substrate to 0.0070, consistent with the pronounced improvement observed in the heatmap analysis. For esterases (Fig. 3d), both models assigned near-unity scores to the true substrate while suppressing the structurally similar non-substrate to near zero (substrate scores of 0.9948 and 0.9883, non-substrate scores of 0.0065 and 0.0028), consistent with the robust margins observed across all similarity bins. For phosphatases (Fig. 3e), fine-tuning elevated the substrate score from 0.8374 to 0.9995. A similar rescue effect was observed for glycosyltransferases (Fig. 3f), where the pretrained model yielded a substrate score of only 0.4040, insufficient for confident classification, whereas fine-tuning recovered it to 0.8709 while maintaining a low non-substrate score (0.0159). DUFs are excluded from panels c–f because their functional annotations are insufficiently characterized to select representative substrate–non-substrate pairs. Together, these analyses demonstrate that fine-tuning primarily benefits families whose substrate chemistries are underrepresented in the pretraining corpus, while families with abundant training coverage already achieve strong discrimination from the pretrained model alone.

### 2.4 Co-folding captures ligand-induced conformational changes inaccessible to rigid docking

As demonstrated by the performance degradation observed when substituting co-folded structures with rigid docking structures in our ablation studies (Section 2.2), high-resolution structural modeling of the bound state represents the cornerstone of accurate enzyme–substrate specificity prediction. To provide detailed mechanistic insight, we examined pig heart citrate synthase, a classic model of large-scale ligand-dependent domain closure in enzyme catalysis. Upon substrate binding, the small domain undergoes an approximately 19-degree rotation relative to the large domain, repositioning three key catalytic residues, His274, His320, and Asp375, into the geometry required for the condensation of acetyl-CoA with oxaloacetate [41, 42]. We compared three distinct structural configurations against the product-bound closed crystal structure (PDB 2CTS, in complex with CoA and citrate) as the reference for the catalytically competent conformation (Fig. 4). These models integrated into our evaluation comprised the unbounded apo crystal structure (PDB 3ENJ), a conventional rigid-body docking pose of acetyl-CoA constructed from an AlphaFold2-predicted structure with an AlphaFill-identified pocket [20, 43], and the Boltz-2 co-folded complex with acetyl-CoA. Because acetyl-CoA and its product CoA-SH differ exclusively by a thioester-linked acetyl group, and the product remains trapped within the active site pocket prior to domain reopening, the closed 2CTS structure provides an experimentally validated reference for the catalytically competent closed conformation.

**Fig. 4.**
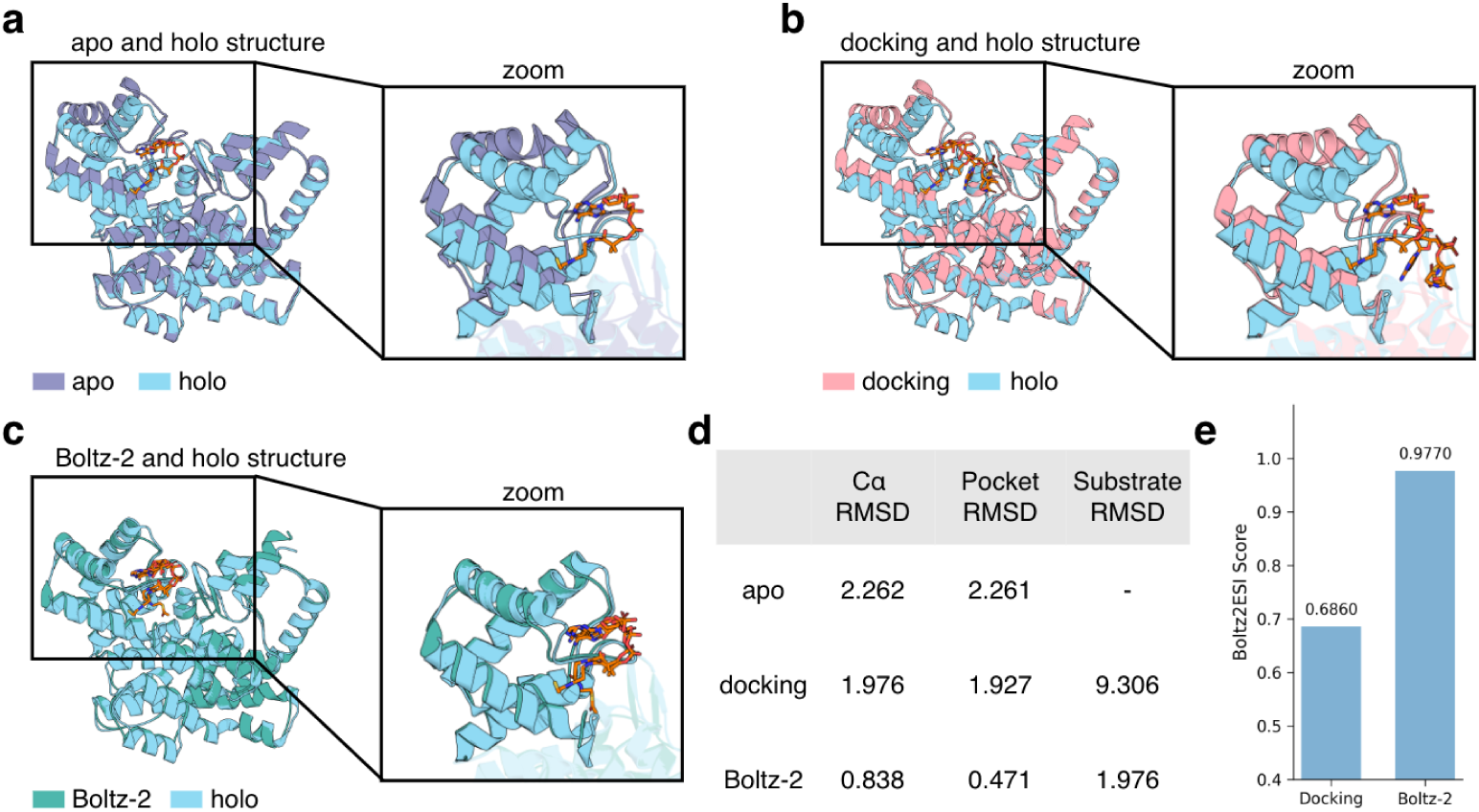
Structural comparison of apo, product-bound closed, rigid docking, and co-folded complexes for pig heart citrate synthase. Apo (PDB 3ENJ) and product-bound closed (PDB 2CTS) crystal structures serve as structural references, rigid docking and co-folding both use acetyl-CoA as input. a, Overlay of the apo crystal structure (PDB 3ENJ; purple) and the product-bound closed crystal structure (PDB 2CTS; blue), with active-site zoom. b, Overlay of the rigid-body docking pose of acetyl-CoA (pink) and the 2CTS reference (blue). c, Overlay of the Boltz-2 co-folded complex (green) with acetyl-CoA (teal) and the 2CTS reference. d, Quantitative RMSD comparison of the three structures relative to the 2CTS structures. e, Boltz2ESI interaction scores for the rigid docking and Boltz-2 co-folded complexes, evaluated on a checkpoint from which pig heart citrate synthase is held out.

We defined the binding pocket as all protein residues with any heavy atom within 5 Å of any holo ligand heavy atom in 2CTS, and computed pocket RMSD as the C*α* RMSD of these residues after Kabsch alignment. Ligand RMSD was obtained by applying the same pocket-derived transform and using Hungarian matching by element type on ligand heavy atoms. Structural alignment reveals that the native apo and product-bound closed configurations differ fundamentally in active-site geometry, exhibiting a global C*α* RMSD of 2.262 Å and a localized pocket RMSD of 2.261 Å. This pronounced domain closure is characteristic of the catalytically competent state of citrate synthase, indicating that the closed conformation is not the predominant state in the absence of substrate and cannot be anticipated by static, single-chain structure prediction alone. When executing conventional rigid-body docking, where the substrate molecule is computationally fitted into a frozen protein scaffold, the pipeline fails to simulate this essential backbone adjustment because the protein backbone is treated as a rigid entity. To quantify the downstream functional impact of this structural divergence, we evaluated both the rigid docking and co-folded configurations using our interaction module. Specifically, we employed the checkpoint trained on the second fold of the unknown substrate and enzyme split, in which all datapoints containing pig heart citrate synthase (UniProt P00889) are held out from the training set. The unadapted, static geometry of the AlphaFill-identified pocket yielded a compromised Boltz2ESI score of 0.6860, whereas the end-to-end co-folded active site recovered an optimal structural fitness, generating a robust catalytic specificity score of 0.9770 (Fig. 4e).

This case study illustrates the biophysical limitations inherent to decoupled, rigid-body docking workflows and underscores how co-folding architectures address them. When utilizing static structural templates, the unadapted pocket configuration alters the topological feature representation provided to the deep-learning module, leading to a degraded capacity to isolate true positive substrates from decoy compounds. This outcome demonstrates that post-hoc docking into rigid scaffolds inherently struggles to capture the coordinated backbone rearrangements and side-chain adjustments required for true enzyme-substrate complementarity [25, 26]. In contrast, the end-to-end co-folding paradigm addresses this bottleneck by optimizing the protein and small molecule coordinates simultaneously, natively capturing the ligand-induced structural plasticity and large-scale domain rearrangements essential for transition-state competence. By explicitly modeling these dynamic structural adaptations rather than relying on unaligned static templates, the co-folding architecture supplies the downstream interaction module with a high-fidelity physical feature space, directly manifested in its superior predictive metrics. Consequently, this capacity to resolve flexible active-site environments provides a robust biophysical explanation for our framework’s superior discriminative resolution, establishing end-to-end co-folding as an essential architectural paradigm for predicting specificity across plastic enzyme superfamilies.

### 2.5 Pathway reconstruction in Withanolide biosynthesis

To evaluate whether Boltz2ESI generalizes beyond its training benchmark, we applied it to a real-world biosynthetic pathway discovery task not represented in the training set. We conducted a retrospective external validation using recently discovered withanolide biosynthetic pathway. Withanolides are phytosterol lactones belonging to the class of triterpenoids and represent the primary bioactive metabolites produced by *Withania somnifera*, commonly known as ashwagandha. Extracts of this plant have been widely used as dietary supplements for improving stress and sleep [44, 45]. Its main bioactive compounds, the withanolides, have been reported to exhibit pharmacological properties such as anti-inflammatory, neuroprotective, and anticancer activities [46]. Recently, Reynolds et al. identified the biosynthetic pathway leading to withanolide intermediates that share chemical features common to withanolide derivatives, including an enone group in ring A and a lactone ring in the side chain [47]. Several cytochrome P450 enzymes, particularly CYP87G1, CYP88C7, and CYP749B2, catalyze the key oxidative reactions in early-stage pathway (Fig. 5a).

**Fig. 5.**
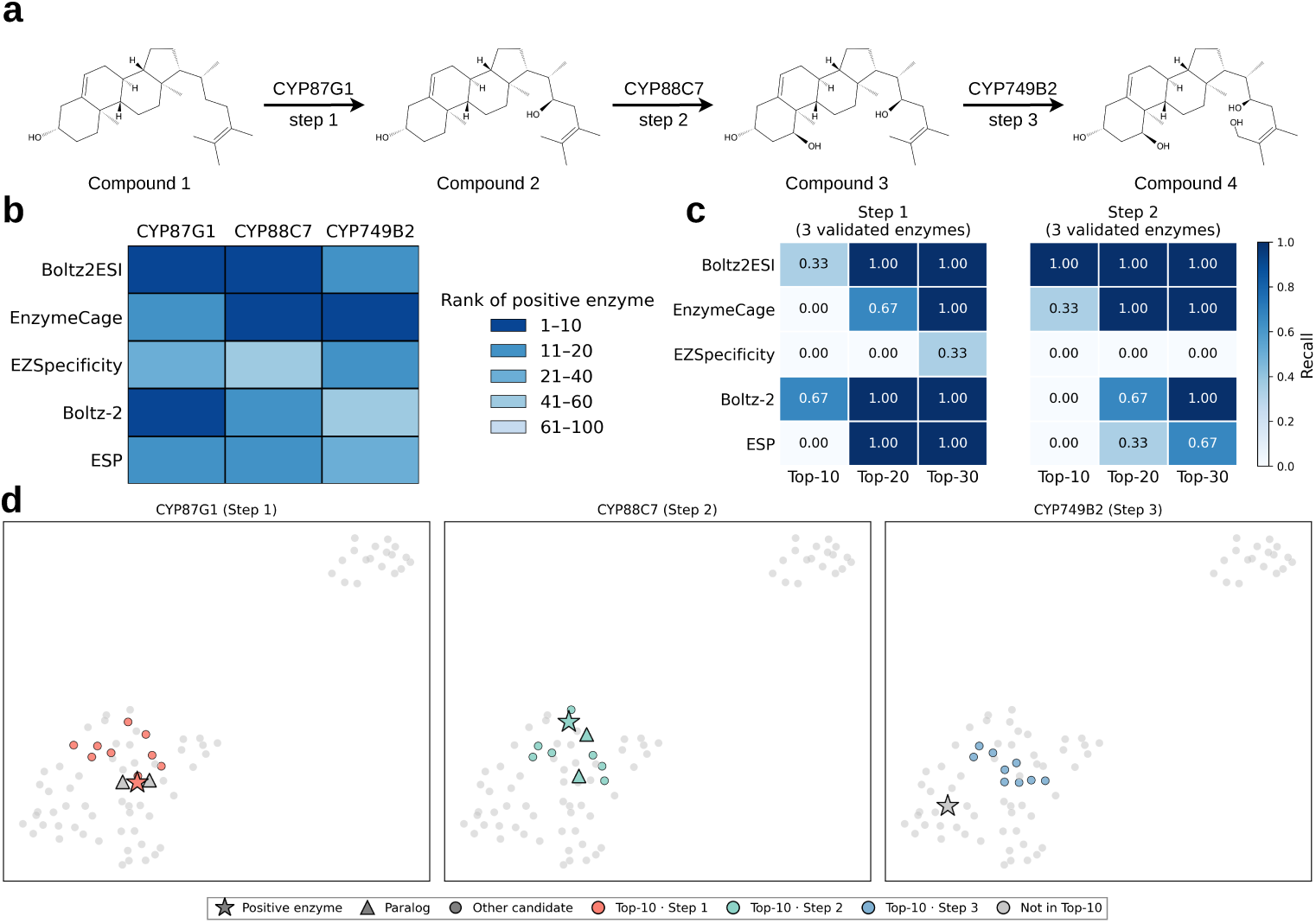
Boltz2ESI prioritizes catalytic CYP enzymes in the Withanolide biosynthetic pathway. **a,** Three-step oxidative cascade in Withanolide biosynthesis, mediated by CYP87G1 (step 1), CYP88C7 (step 2), and CYP749B2 (step 3). **b,** Rank of each confirmed CYP enzyme among candidate plant P450 sequences, for Boltz2ESI and four baselines. Color encodes rank range. **c,** Recall at top-*k* (*k* = 10, 20, 30) for recovering the true CYP and its biochemically characterized paralogs (two paralogs for steps 1 and 2, respectively) from the candidate pool. **d,** UMAP visualization of the 107 P450 candidates based on Foldseek structural tokens and ESM3 sequence representations, shown separately for each biosynthetic step. Stars denote validated positive enzymes, triangles denote biochemically characterized paralogs, and colored circles denote Boltz2ESI top-10 ranked candidates.

To evaluate the model’s ability to recover experimentally validated P450 enzymes involved in the biosynthetic transitions of withanolide intermediate compounds (Fig. 5a), we applied Boltz2ESI to a candidate pool of 107 P450 sequences identified from a *de novo* assembled *W. somnifera* transcriptome via Pfam domain annotation. This candidate pool was also adopted by EnzymeCAGE [4], enabling a direct one-to-one comparison. We then scored all pairwise interactions between these 107 candidates and the three withanolide intermediates using Boltz2ESI, without any pathway-specific training or fine-tuning. Boltz2ESI ranked CYP87G1 8th and CYP88C7 4th among the 107 candidates, outperforming EnzymeCAGE (13th and 10th, respectively; Fig. 5b, Supplementary Table 6). For step 3, EnzymeCAGE ranked CYP749B2 6th while Boltz2ESI ranked it 17th. One possible explanation is that CYP749B2 catalyzes a more complex transformation involving multiple oxidative steps followed by spontaneous lactone ring formation, producing a product structurally more divergent from the preceding intermediates. Such multi-step transformations may be underrepresented in current training data.

To assess broader functional coverage, we expanded the evaluation to include paralogs of the enzymes responsible for steps 1 and 2. Specifically, we added two CYP87G1 and two CYP88C7 paralogs, which have been experimentally shown to catalyze the same reactions as the reference enzymes, albeit with equal or lower catalytic activity [47], to the candidate pool, rescored all candidates, and computed recall at top-*k* (*k* = 10, 20, 30; Fig. 5c).

For CYP87G1, the additional paralogs CYP87G1-1 and CYP87G1-3 share 94% and 99% amino acid sequence identity with CYP87G1-2, respectively, the latter exhibiting the highest catalytic activity. Despite their high sequence similarity, these paralogs display distinct tissue-specific expression patterns: CYP87G1-1 is predominantly expressed in roots, whereas CYP87G1-2 and CYP87G1-3 are mainly expressed in leaves. [47] Boltz2ESI ranked CYP87G1 paralogs 12th, 10th, and 13th for CYP87G1-1, CYP87G1-2, and CYP87G1-3, respectively. It not only achieved the highest recall, with perfect top-20 recall (recall = 1.00), but also showed improved localization of CYP87G1 paralogs within the ranking, indicating more coherent clustering (Supplementary Table 6).

Similarly, for CYP88C7, the additional paralogs CYP88C7-2 and CYP88C7-3 share 99.8% and 90% amino acid sequence identity, respectively, with CYP88C7-1, which exhibits the highest catalytic activity among the functional paralogs. Similar tissue-specific expression patterns were observed, with CYP88C7-1 and CYP88C7-2 predominantly expressed in leaves, and CYP88C7-3 mainly in roots [47]. Boltz2ESI ranked CYP88C7 paralogs 5th, 4th, and 7th for CYP88C7-1, CYP88C7-2, and CYP88C7-3, respectively. It also achieved the highest recall, with perfect top-10 recall (recall = 1.00), indicating more coherent clustering for step 2 as well (Supplementary Table 6).

Reynolds et al. [47] further reported that these paralogs exhibited tissue-specific expression patterns, prompting investigation of potential functional divergence between root- and leaf-expressed enzymes, which was later assessed in an orthologous host system where no differences in catalytic activity were observed among the paralogs. Consistently, Boltz2ESI correctly grouped all paralogs within the ranking, in agreement with their shared catalytic behavior. This indicates that Boltz2ESI could potentially capture underlying molecular function directly from structural features, independent of gene expression context, providing a structural perspective that complements gene expression–based omics approaches.

To further examine what the model captures, we visualized the candidates and their paralogs using UMAP projections derived from Foldseek structural tokens and ESM3 sequence representations (Fig. 5d). Across all three biosynthetic steps, the Boltz2ESI top-10 ranked candidates cluster in proximity to the validated positive enzyme, and the biochemically characterized paralogs co-localize with the positive enzyme in the embedding space. The consistency between this structural–evolutionary embedding space and the Boltz2ESI prioritization provides qualitative support that the model preserves biologically meaningful relationships among homologous enzymes.

Taken together, these results demonstrate that Boltz2ESI generalizes effectively to secondary metabolite biosynthetic enzymes phylogenetically distant from its training data, supporting its utility for candidate prioritization in natural product pathway elucidation.

## 3 Discussion

Enzyme–substrate specificity is ultimately encoded in the three-dimensional interplay between a flexible protein scaffold and its cognate small molecule. Boltz2ESI leverages this insight by extracting representations directly from a co-folding foundation model that jointly optimizes enzyme and substrate coordinates, capturing the ligand-induced conformational plasticity that rigid-body docking pipelines are structurally unable to model. The citrate synthase case study (Fig. 4) provides a concrete illustration: only co-folding recovers the domain closure essential for catalytic competence, whereas conventional docking into a frozen apo scaffold leaves the substrate severely mispositioned. Because the resulting pairwise and single representations encode enzyme–substrate interaction geometry at a resolution finer than sequence identity, Boltz2ESI discriminates among enzymes sharing the same four-digit EC number (Fig. 2b). Moreover, co-folding localizes the active site as part of the structure prediction itself, eliminating the dependence on predefined binding-pocket annotations and extending applicability to orphan enzymes whose active sites have not been structurally catalogued.

A key finding from our ablation study is that high-fidelity structural modeling and evolutionary context play complementary rather than redundant roles (Fig. 2c). Removing ESM3 embeddings produced the largest single-component performance drop, indicating that global evolutionary context remains indispensable for generalizing to sequence-distant enzymes. At the same time, replacing co-folded complexes with rigid docking poses caused the most severe AUPR degradation, confirming that the quality of the structural complex is the primary bottleneck for precision. Family-level evaluation shows that this multi-modal foundation generalizes broadly. The ESIBank-pretrained model exceeds baselines on most families without modification, while lightweight fine-tuning on modest family-specific data recovers performance for underrepresented families such as thiolases and DUFs (Fig. 3). A novelty analysis stratifying test-set performance by the maximum sequence identity to training enzymes and the maximum Tanimoto similarity to training substrates further showed that Boltz2ESI retained discriminative performance across enzyme and substrate similarity regimes, including samples with low similarity to the training data (Supplementary Figs. 3 and 4).

The Withanolide pathway reconstruction highlights both the strengths and the boundaries of our approach. Boltz2ESI outperformed EnzymeCAGE on the first two biosynthetic steps, where the catalytic CYPs are phylogenetically distant from known structural templates, but ranked CYP749B2 lower on step 3. This discrepancy likely reflects the higher sequence similarity between CYP749B2 and structurally characterized entries in EnzymeCAGE’s training corpus, a regime where template-based methods can directly transfer geometric information without de novo structure generation. Co-folding and homology-guided approaches are therefore complementary: co-folding provides the greater advantage for conformationally plastic or phylogenetically isolated enzymes, while homology transfer remains effective when high-quality structural templates already exist. Beyond individual enzymatic steps, this study suggests implications for biosynthetic pathway reconstruction beyond omics-based approaches. Omics methods infer functional relationships from gene expression data, often assuming correlated expression among pathway genes [7]. However, this assumption can be confounded by context-dependent temporal and spatial regulation. In contrast, Boltz2ESI leverages structural information to predict protein–ligand interactions based on intrinsic molecular features rather than expression signals. These approaches are complementary, where omics captures system-level context, while structure-based models provide mechanistic insight at the molecular level. Integrating both offers a more robust and biologically grounded strategy for pathway elucidation.

Our evaluation also inherits limitations of the underlying data. The ESIBank benchmark constructs negatives by pairing enzymes with substrates from related EC classes, yet BRENDA annotations are incomplete: some nominally negative pairs may in fact be catalytically competent (pseudo-negatives), biasing measured performance downward for all methods and inflating the apparent difficulty of the benchmark. Similarly, functional annotations for DUFs and other poorly characterized families carry inherent uncertainty, and experimental validation remains essential before drawing mechanistic conclusions from model predictions in these classes.

From a practical standpoint, the two-stage co-folding protocol is computationally more demanding than sequence-only or docking-based pipelines, which may limit throughput for proteome-scale screening. Distilling the co-folding representations into lighter architectures, or amortizing inference by caching enzyme structures across multiple substrate queries, are promising directions for scaling. More broadly, as co-folding models continue to improve in accuracy and efficiency, we anticipate that the framework introduced here can be extended to cofactor specificity prediction, enzyme–inhibitor discrimination, and de novo pathway design.

## 4 Methods

### 4.1 ESIBank benchmark

We trained and evaluated Boltz2ESI on the ESIBank dataset [19], a comprehensive enzyme–substrate interaction benchmark spanning 34,417 unique substrates and 8,124 unique enzymes (323,783 high-quality pairs after quality filtering). The BRENDA-derived core was constructed by collecting native and non-native substrate SMILES from BRENDA [33] and matching each entry to a representative enzyme sequence in UniProt [6] via EC number and organism annotation. For each positive pair, five negative enzymes and five negative substrates were sampled using the EC number hierarchy, producing ten negative pairs that span five difficulty levels, from zero shared EC digits to all four digits identical, so that the resulting negatives range from trivially distinguishable to highly challenging decoys. Quality filters removed substrates containing more than 280 heavy atoms, enzymes exceeding 1,000 residues, and entries lacking annotated active-site information in UniProt. The BRENDA core was supplemented with curated family-specific datasets for six representative enzyme families: thiolases [34], esterases [35], phosphatases [36], glycosyltransferases [37], nitrilases [38] and domain of unknown function proteins (DUFs) [39]. We adopted two data split regimes from EZSpecificity [19]. In the random split, enzyme–substrate pairs are partitioned independently of identity. In the unknown substrate and enzyme split, all pairs containing a given enzyme or substrate are assigned to the same fold, ensuring that neither the enzyme sequence nor the substrate appears in both training and test sets. All experiments used four-fold cross-validation (split statistics in Supplementary Tables 2 and 3).

### 4.2 Enzyme–substrate co-folding and structural representations

Existing structure-aware methods [4, 19] depend on predefined binding pockets, typically transferred from homologous structures via AlphaFill [43], and employ rigid-body docking to position the substrate, precluding faithful modeling of the mutual adaptation between the substrate and its binding environment. Boltz2ESI overcomes these limitations by employing two-stage end-to-end co-folding for enzyme-substrate pairs, using Boltz-2 [27] to jointly predict the complex structure and extract its internal representations.

In the first stage, a full-length co-folding run is performed with multiple sequence alignments (MSAs) to produce five candidate complex structures, from which the highest-confidence prediction is selected. This global structure serves to localize the putative active site. Following Boltz-2 affinity module, the active site extractor then retains all substrate heavy atoms and selects up to 200 neighbouring protein residues by proximity (with sequential context around each selected residue), yielding a focused active-site pocket [27]. In the second stage, the cropped active-site pocket and the substrate are re-folded by Boltz-2 without MSA input. Five refined pocket–ligand structures are generated, and the structure with the highest interface predicted TM-score (ipTM) is selected.

From the selected active-site complex, we extract three complementary structural representations: (i) the per-token single representation **s**^input^ ∈ ℝ*^N×d_s_^*, encoding chemical identity for each amino acid residue and substrate heavy atom; (ii) the pairwise trunk representation **z**^trunk^ ∈ ℝ*^N×N×d_z_^*, encoding active site residues and substrate atom interaction geometry learned across Boltz-2’s large-scale co-folding pretraining; and (iii) a distogram **d**^complex^ ∈ ℝ*^N×N×d_d_^*, discretizing the pairwise distance distribution of the predicted complex. In all three representations, each token corresponds to either an amino acid residue in the active site or a heavy atom in the substrate.

### 4.3 Evolutionary and molecular priors extraction

#### Enzyme evolutionary representations

Since the second-stage co-folding operates on the cropped active-site pocket without MSA, the resulting structural representations lack global evolutionary context. To complement this, we first encode the full-length enzyme sequence with ESM3-1.4B [28], obtaining per-residue embeddings, and then extract the subset corresponding to the active-site residues. These embeddings capture evolutionary conservation, protein stability signals and distal allosteric effects that are not accessible from the local pocket structure alone.

#### Substrate molecular representations

Substrate chemistry is encoded through two complementary descriptors. Uni-Mol2 [29] (84M parameters) encodes the substrate three-dimensional conformation, generated from SMILES using RDKit [48], into a 512-dimensional molecular embedding via geometry-aware pretraining. Morgan fingerprints [30] with a radius of 2 and 2,048 bits encode topological substructure information, providing a complementary representation of molecular identity and functional group composition. The Uni-Mol2 embedding and Morgan fingerprint are concatenated to form a unified ligand feature representation.

### 4.4 Enzyme–substrate interaction prediction module

The interaction prediction module fuses the co-folding structural representations with the evolutionary and molecular priors to produce a catalytic interaction probability.

#### Single-representation fusion

The co-folding single representation **s**^input^ is fused with the external priors through separate projections for enzyme and substrate tokens. For enzyme residues, the ESM3 embeddings **s**^esm^ are concatenated with **s**^input^ and projected by a two-layer multilayer perceptron (MLP) head:

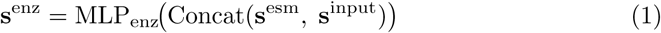

For substrate atoms, the ligand feature vector **f**^lig^ (Uni-Mol2 embedding concatenated with Morgan fingerprint) is similarly fused:

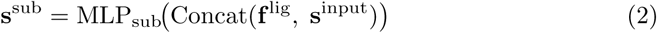

where MLP_enz_ has a hidden dimension of 1,024 with dropout 0.1 and MLP_sub_ has a hidden dimension of 384. Both of them use GELU activations and output 384-dimensional representations.

#### Pair-representation conditioning

We followed the Boltz-2 affinity module [27], the trunk pair representation **z**^trunk^ is masked to zero out all protein–protein entries, restricting the representation to protein–ligand and ligand–ligand pairs. The fused single representations are then broadcast into the pair dimension, and the distogram **d**^complex^ is added as a pairwise conditioning signal.

#### Pairformer and output head

The conditioned pair tensor is processed through a four-block Pairformer stack [31], in which attention is likewise masked to exclude protein–protein pairs, so that message passing is restricted to cross-modal interactions. The Pairformer is initialized with pre-trained weights from the Boltz-2 affinity head. The output pair representation is reduced via mean pooling over all protein–ligand and ligand–ligand token pairs, and a multilayer perceptron head maps the pooled representation to a scalar logit. The final interaction probability is obtained as:

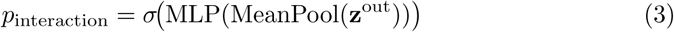

where *σ* denotes the sigmoid function. The complete pipeline pseudocode and architectural hyperparameters are provided in Supplementary Note 2 and Supplementary Table 1.

### 4.5 Training procedure

Co-folding structures, ESM3 embeddings, Uni-Mol2 embeddings and Morgan fingerprints were pre-computed offline for all enzyme–substrate pairs. During training, only the parameters of the interaction prediction module were updated. The model was trained with binary cross-entropy loss and optimized using AdamW with a learning rate of 1 × 10*^−^*^4^, weight decay of 1 × 10*^−^*^5^ and gradient clipping at a maximum norm of 1.0. We applied a reduce-on-plateau learning rate schedule that monitored validation AUROC. When AUROC did not improve for 5 consecutive epochs, the learning rate was multiplied by a factor of 0.5. Training ran for a maximum of 15 epochs. We used a batch size of 64 across eight NVIDIA L40S GPUs in FP32 precision. Each cross-validation fold required approximately 35 hours. For final evaluation, within each cross-validation fold we retained two checkpoints, the one with the highest validation AUROC and the one with the highest validation AUPR. The reported metrics are the average of these two checkpoints across all four folds. For family-specific fine-tuning (Boltz2ESI-fine-tune), we started from the ESIBank-pretrained checkpoint with the batch size reduced to 8 on a single GPU. The learning rate schedule monitored validation AUPR, and we selected the single checkpoint with the highest validation AUPR for evaluation. Software and computational environment are listed in Supplementary Note 3.

## Acknowledgements

This work was supported in part by the NVIDIA Academic Grant Program and the Google TPU Research Cloud. W.J. acknowledges support from the Google Research Scholar Award, the Torrey Coast Foundation, and the Defense Advanced Research Projects Agency (DARPA). W.Y.K. acknowledges support from the National Research Foundation of Korea (NRF), funded by the Ministry of Science and ICT (MSIT) of the Korean government (grant no. NRF-2023R1A2C2004376).

## Supplementary Information

## Supplementary Note 1: Baseline implementation details

We compared Boltz2ESI with the following baseline methods. Below we describe each model and the details of how it was applied in our evaluation.

### ESP

ESP [40] is a machine learning model for predicting enzyme–substrate pairs across diverse enzyme families. It uses a modified ESM-1b transformer to generate protein embeddings and GNN-based fingerprints to capture substrate features. These representations are combined and fed into a gradient-boosted decision tree to predict enzyme–substrate relationships. ESP takes enzyme–substrate pairs as input, which is directly compatible with the ESIBank format. Because ESP was trained on a different dataset, we used the officially released model weights and inference scripts to score all enzyme–substrate pairs in the ESIBank benchmark and Withanolide pathway evaluation.

### Boltz-2

Boltz-2 [27] is a biomolecular foundation model capable of predicting binding complex structures and estimating binding affinities. Its official release includes two affinity head checkpoints, the default inference pipeline ensembles predictions from both heads to produce a final affinity score. We evaluated Boltz-2 as a zero-shot baseline by running the official affinity prediction pipeline on the co-folded enzyme–substrate complexes. The predicted binding affinity was used directly as the interaction score without any task-specific training or fine-tuning. Boltz-2 takes enzyme–substrate pairs as input, which directly aligns with the ESIBank data format without further conversion.

### EZSpecificity

EZSpecificity [19] is a cross-attention graph neural network that predicts enzyme– substrate specificity by jointly encoding enzyme active-site structures and substrate molecular graphs. The model takes enzyme–substrate pairs as input, matching the format of the ESIBank benchmark. Because the authors of EZSpecificity constructed the ESIBank benchmark and trained their model on it, we directly adopted their published benchmark results for the ESIBank evaluation to ensure a fair comparison under identical data splits and cross-validation folds. We note that EZSpecificity only open-sourced model weights for the random split; results on the unknown substrate and enzyme split are therefore taken from the original publication. For the Withanolide pathway evaluation, both EZSpecificity and Boltz2ESI used models trained on the random split, as this external test set is independent of the ESIBank training data.

### EnzymeCAGE

EnzymeCAGE [4] is a geometric foundation model for predicting enzyme–reaction compatibility. It encodes enzyme pocket structures using a geometric vector perceptron (GVP) network and reaction representations using a SchNet-based 3D molecular encoder, then fuses the two modalities through a geometry-enhanced cross-attention interaction module to produce a compatibility score. The model supports both enzyme retrieval (given a reaction, rank candidate enzymes) and reaction retrieval (given an enzyme, rank candidate reactions). Unlike the methods above, EnzymeCAGE takes enzyme–reaction pairs rather than enzyme–substrate pairs as input. We evaluated EnzymeCAGE exclusively on the Withanolide biosynthetic pathway test set, where reaction annotations are available. We used the officially released model and inference code to score all candidates against the positive enzymes.

## Supplementary Note 2: Boltz2ESI architecture and hyperparameters

The complete Boltz2ESI pipeline, the active-site extraction procedure, and the enzyme–substrate interaction (ESI) prediction module are described in Algorithms 1, 2, and 3, respectively. Supplementary Table S1 summarises the key architectural and inference hyperparameters not specified in the main text.

### Algorithm 1

**Boltz2ESI pipeline**

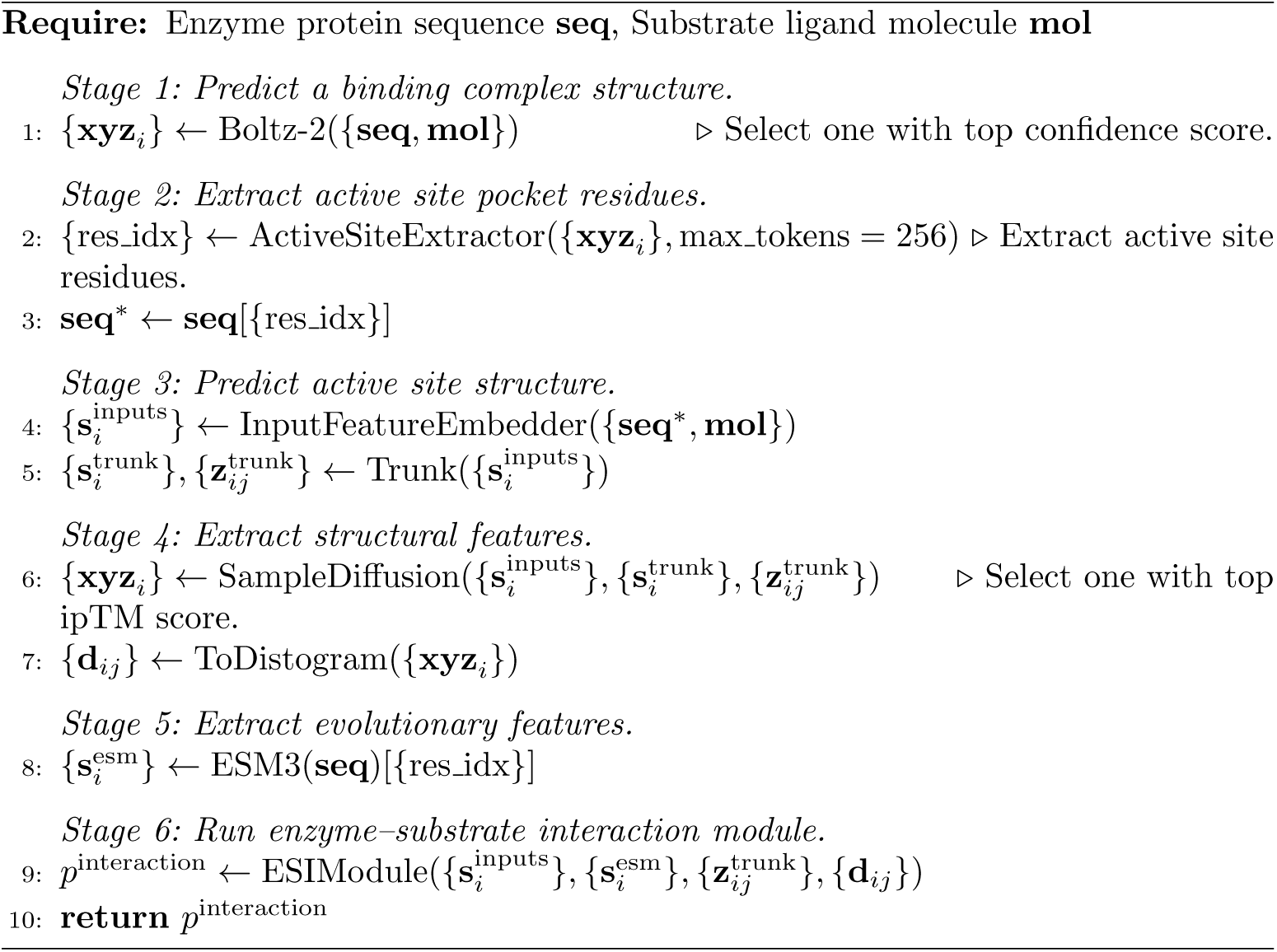

### Algorithm 2

**Active–site extraction**

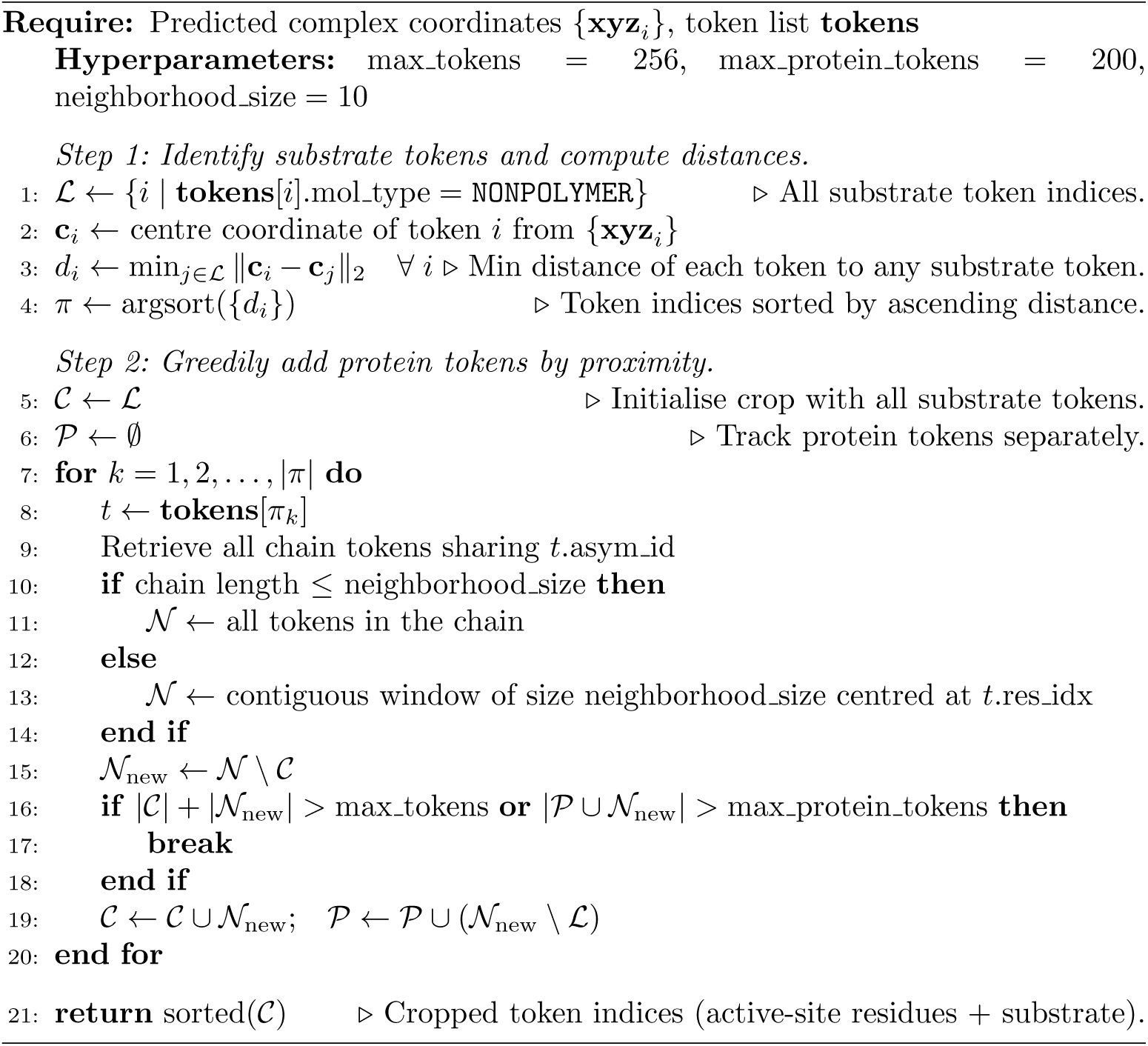

### Algorithm 3

**Enzyme–substrate interaction prediction module**

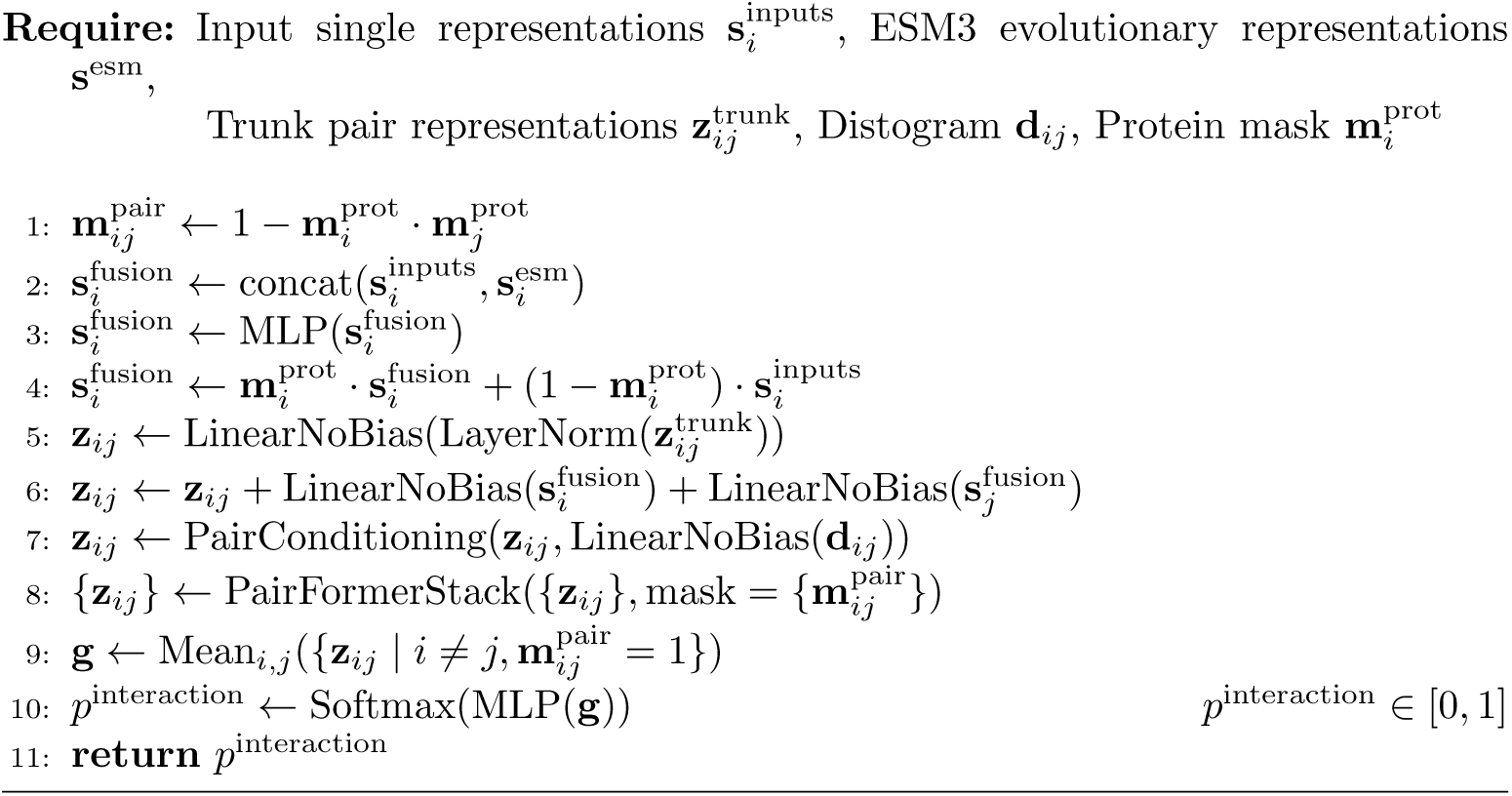

## Supplementary Note 3: Software and computational environment

All experiments were conducted on a single node equipped with 8× NVIDIA L40S GPUs (46 GB VRAM each). The following software versions were used throughout the experiments:

- Boltz-2 v2.2.1 (commit 8ed3100)
- ESM3: esm v3.2.3 (esm3 sm open v1)
- Foldseek v10.941cd33 (for structural analysis)

## Supplementary Tables

**Table S1.** Architectural and inference hyperparameters of Boltz2ESI.

| Parameter | Value |
| --- | --- |
| PairFormer layers | 4 |
| Max pocket residues | 200 |
| Num recycles (full complex) | 3 |
| Num diffusion samples (full complex) | 5 |
| Num recycles (active site) | 5 |
| Num diffusion samples (active site) | 5 |

**Table S2.** Total positive and negative case counts and their train, validation, and test set distribution under four-fold cross-validation for six enzyme families of the random split in ESIBank.

| Family | Class | Total | Split 1 |  |  | Split 2 |  |  | Split 3 |  |  | Split 4 |  |  |
| --- | --- | --- | --- | --- | --- | --- | --- | --- | --- | --- | --- | --- | --- | --- |
|  |  |  | Train | Val | Test | Train | Val | Test | Train | Val | Test | Train | Val | Test |
| Thiolases | Positive | 550 | 414 | 41 | 95 | 416 | 38 | 96 | 409 | 43 | 98 | 411 | 47 | 92 |
|  | Negative | 545 | 408 | 40 | 97 | 405 | 44 | 96 | 412 | 39 | 94 | 410 | 35 | 100 |
| Esterases | Positive | 3,076 | 2,308 | 236 | 532 | 2,276 | 229 | 571 | 2,350 | 233 | 493 | 2,294 | 253 | 529 |
|  | Negative | 10,940 | 8,204 | 815 | 1,921 | 8,236 | 822 | 1,882 | 8,162 | 818 | 1,960 | 8,218 | 798 | 1,924 |
| Phosphatases | Positive | 5,424 | 4,040 | 406 | 978 | 4,074 | 403 | 947 | 4,051 | 398 | 975 | 4,107 | 400 | 917 |
|  | Negative | 30,546 | 22,938 | 2,291 | 5,317 | 22,903 | 2,294 | 5,349 | 22,927 | 2,299 | 5,320 | 22,870 | 2,297 | 5,379 |
| Glycosyltransferases | Positive | 1,019 | 766 | 74 | 179 | 766 | 69 | 184 | 753 | 86 | 180 | 772 | 67 | 180 |
|  | Negative | 3,995 | 2,995 | 301 | 699 | 2,994 | 307 | 694 | 3,008 | 289 | 698 | 2,988 | 309 | 698 |
| Nitrilases | Positive | 85 | 68 | 3 | 14 | 64 | 6 | 15 | 59 | 7 | 19 | 64 | 6 | 15 |
|  | Negative | 599 | 445 | 48 | 106 | 449 | 45 | 105 | 454 | 44 | 101 | 449 | 45 | 105 |
| DUFs | Positive | 274 | 198 | 21 | 55 | 205 | 17 | 52 | 221 | 14 | 39 | 198 | 21 | 55 |
|  | Negative | 2,463 | 1,855 | 184 | 424 | 1,848 | 188 | 427 | 1,832 | 191 | 440 | 1,854 | 184 | 425 |

**Table S3.** Total positive and negative case counts and their train, validation, and test set distribution under four-fold cross-validation for six enzyme families of the unknown substrate and enzyme split in ESIBank. Nitrilase is excluded from the final evaluation, as certain folds contain no positive samples in the test set.

| Family | Class | Total | Split 1 |  |  | Split 2 |  |  | Split 3 |  |  | Split 4 |  |  |
| --- | --- | --- | --- | --- | --- | --- | --- | --- | --- | --- | --- | --- | --- | --- |
|  |  |  | Train | Val | Test | Train | Val | Test | Train | Val | Test | Train | Val | Test |
| Thiolases | Positive | 550 | 355 | 0 | 12 | 321 | 0 | 26 | 270 | 3 | 27 | 294 | 3 | 20 |
|  | Negative | 545 | 305 | 10 | 1 | 284 | 5 | 13 | 335 | 2 | 12 | 300 | 2 | 22 |
| Esterases | Positive | 3,076 | 1,735 | 15 | 103 | 1,704 | 24 | 95 | 1,848 | 20 | 68 | 1,636 | 20 | 117 |
|  | Negative | 10,940 | 6,185 | 55 | 339 | 6,144 | 53 | 347 | 6,072 | 50 | 374 | 6,212 | 57 | 325 |
| Phosphatases | Positive | 5,424 | 3,152 | 43 | 143 | 3,180 | 20 | 165 | 2,953 | 23 | 223 | 2,954 | 40 | 166 |
|  | Negative | 30,546 | 17,184 | 149 | 959 | 17,032 | 172 | 966 | 17,383 | 169 | 879 | 17,095 | 152 | 1,004 |
| Glycosyltransferases | Positive | 1,019 | 570 | 8 | 35 | 574 | 5 | 36 | 527 | 6 | 37 | 628 | 2 | 26 |
|  | Negative | 3,995 | 2,283 | 12 | 134 | 2,220 | 22 | 127 | 2,270 | 22 | 122 | 2,207 | 19 | 141 |
| Nitrilases | Positive | 85 | 57 | 1 | 2 | 57 | 0 | 0 | 49 | 0 | 5 | 31 | 1 | 9 |
|  | Negative | 599 | 349 | 1 | 19 | 307 | 3 | 28 | 357 | 2 | 16 | 333 | 2 | 19 |
| DUFs | Positive | 274 | 175 | 0 | 7 | 181 | 0 | 7 | 151 | 0 | 13 | 111 | 3 | 18 |
|  | Negative | 2,463 | 1,398 | 12 | 77 | 1,392 | 12 | 77 | 1,422 | 12 | 71 | 1,329 | 9 | 98 |

**Table S4.**
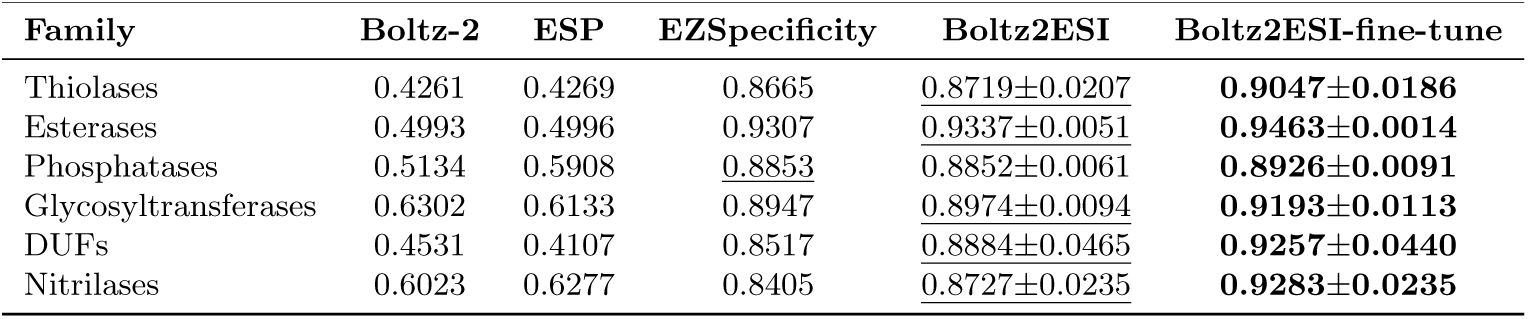
Per-family AUROC on the ESIBank random split. For Boltz2ESI and Boltz2ESI-fine-tune, values are reported as mean *±* standard deviation across four cross-validation folds. The highest and second highest values in each row are indicated by bold and underlined text, respectively.

**Table S5.**
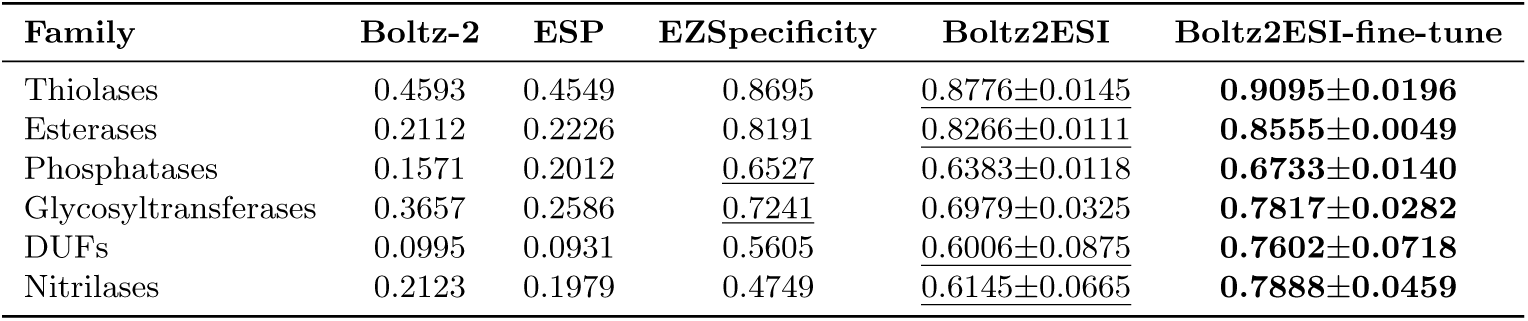
Per-family AUPRC on the ESIBank random split. For Boltz2ESI and Boltz2ESI-fine-tune, values are reported as mean *±* standard deviation across four cross-validation folds. The highest and second highest values in each row are indicated by bold and underlined text, respectively.

**Table S6.**
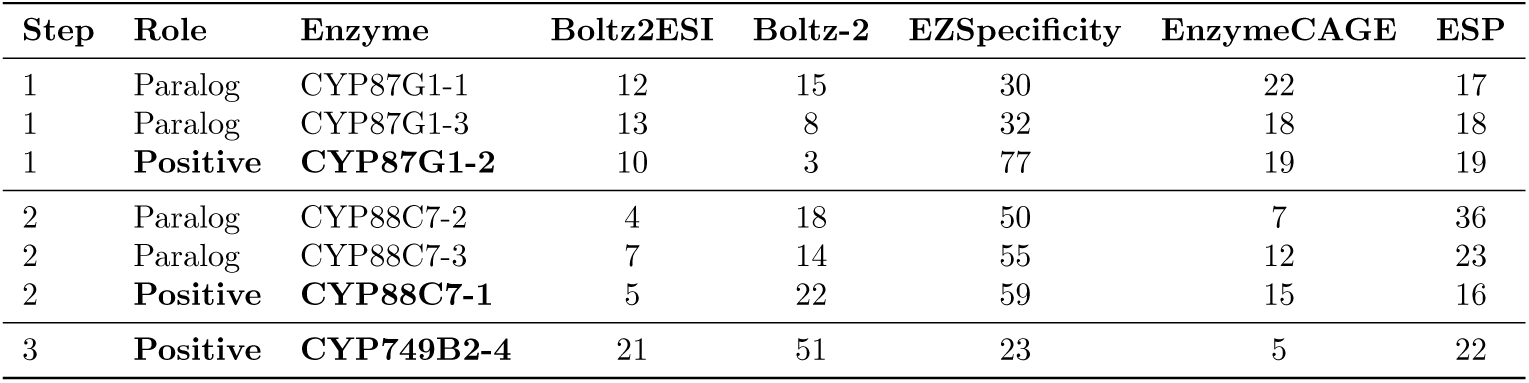
Rank of each validated withanolide CYP enzyme and its biochemically characterized paralogs among candidate P450 sequences for each biosynthetic step.

| Step | Role | Enzyme | Boltz2ESI | Boltz-2 | EZSpecificity | EnzymeCAGE | ESP |
| --- | --- | --- | --- | --- | --- | --- | --- |
| 1 | Paralog | CYP87G1-1 | 12 | 15 | 30 | 22 | 17 |
| 1 | Paralog | CYP87G1-3 | 13 | 8 | 32 | 18 | 18 |
| 1 | **Positive** | **CYP87G1-2** | 10 | 3 | 77 | 19 | 19 |
| 2 | Paralog | CYP88C7-2 | 4 | 18 | 50 | 7 | 36 |
| 2 | Paralog | CYP88C7-3 | 7 | 14 | 55 | 12 | 23 |
| 2 | **Positive** | **CYP88C7-1** | 5 | 22 | 59 | 15 | 16 |
| 3 | **Positive** | **CYP749B2-4** | 21 | 51 | 23 | 5 | 22 |

## Supplementary Figures

**Fig. S1.**
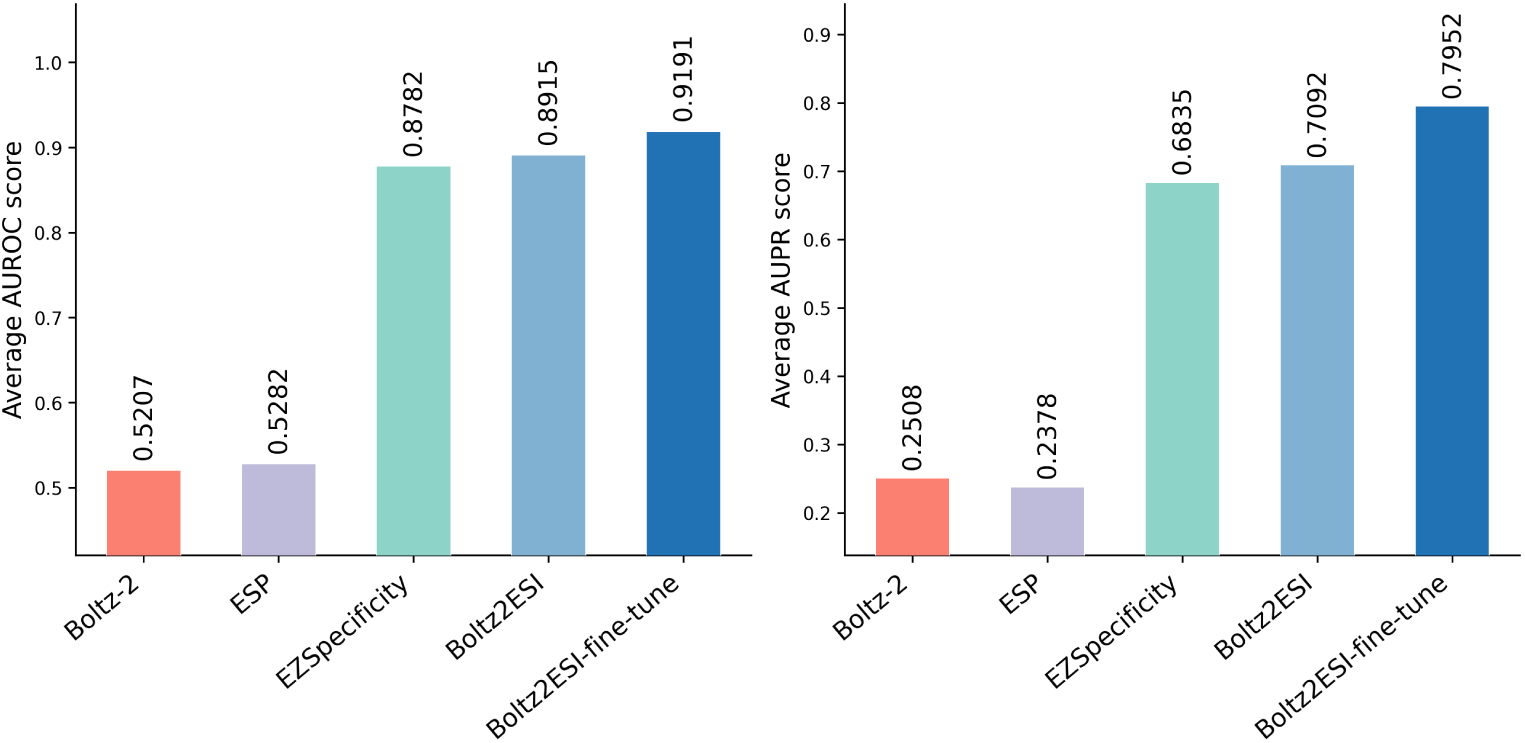
Average per-family AUROC (left) and AUPR (right) on the ESIBank random split, comparing Boltz-2, ESP, EZSpecificity, Boltz2ESI, and Boltz2ESI-fine-tune. Scores are averaged across all six enzyme families.

**Fig. S2.**
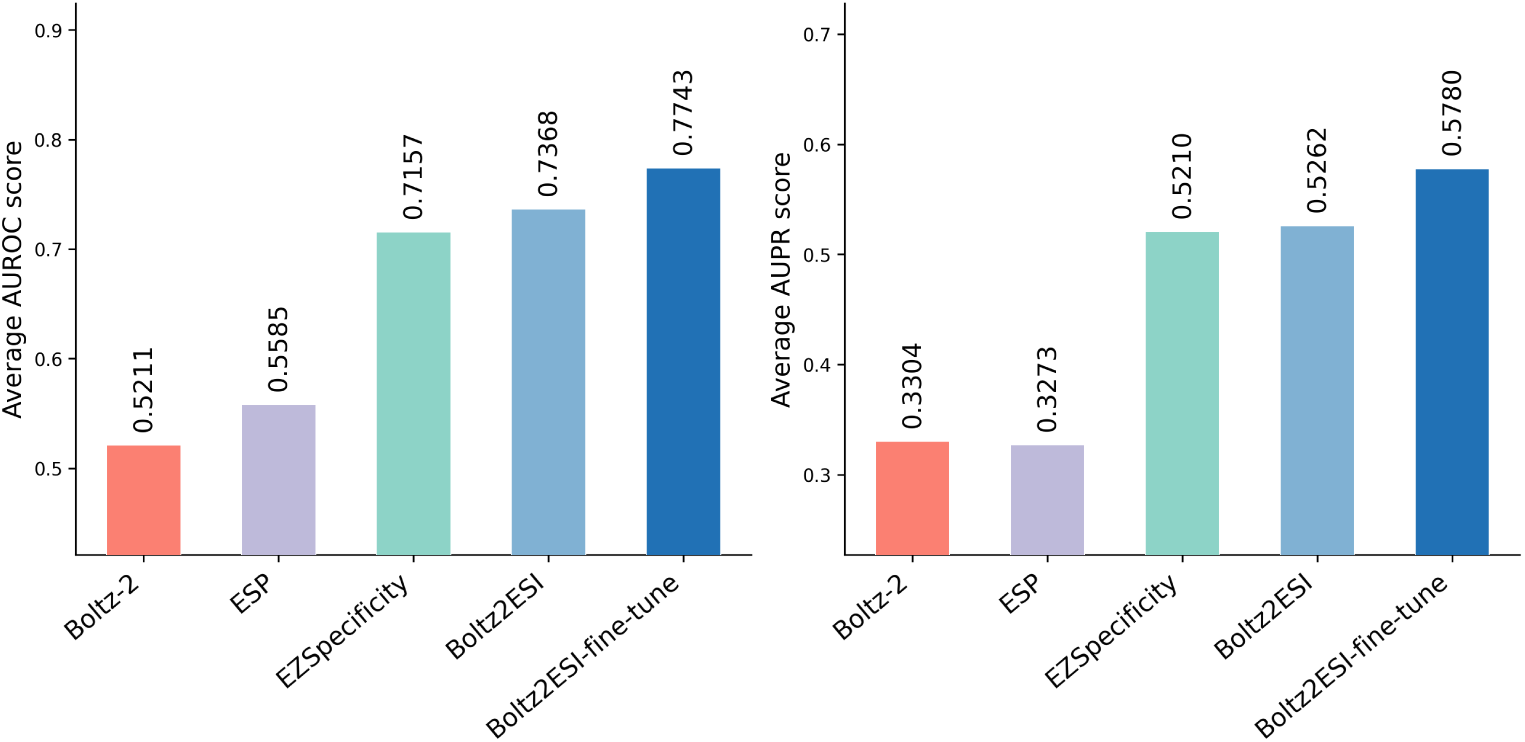
Average per-family AUROC (left) and AUPR (right) on the ESIBank unknown substrate and enzyme split, comparing Boltz-2, ESP, EZSpecificity, Boltz2ESI, and Boltz2ESI-fine-tune. Scores are averaged across five enzyme families (nitrilases excluded owing to insufficient test data).

**Fig. S3.**
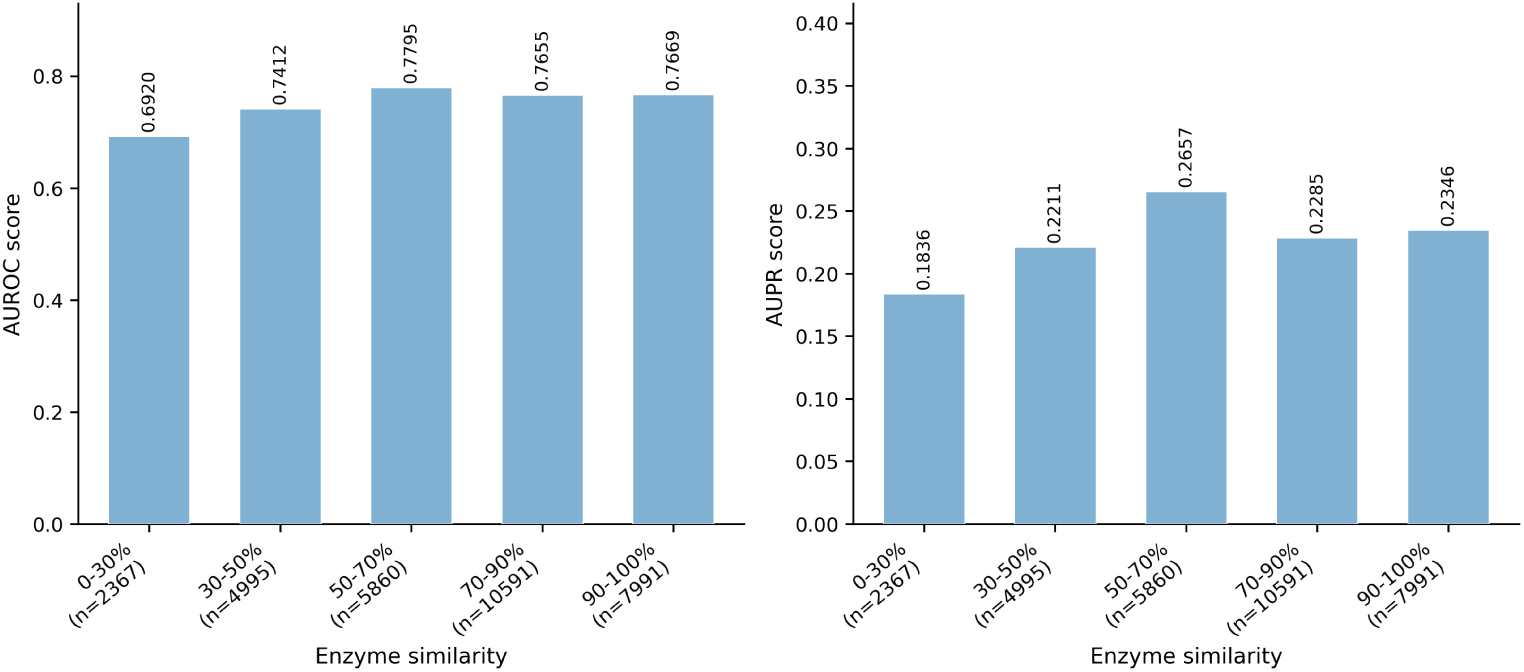
Boltz2ESI performance on the unknown substrate and enzyme split stratified by enzyme novelty. Test-set enzyme–substrate pairs are binned by the maximum MMseqs2 sequence identity between the test enzyme and any training-set enzyme. AUROC (left) and AUPR (right) are shown for each bin, with sample sizes indicated in parentheses.

**Fig. S4.**
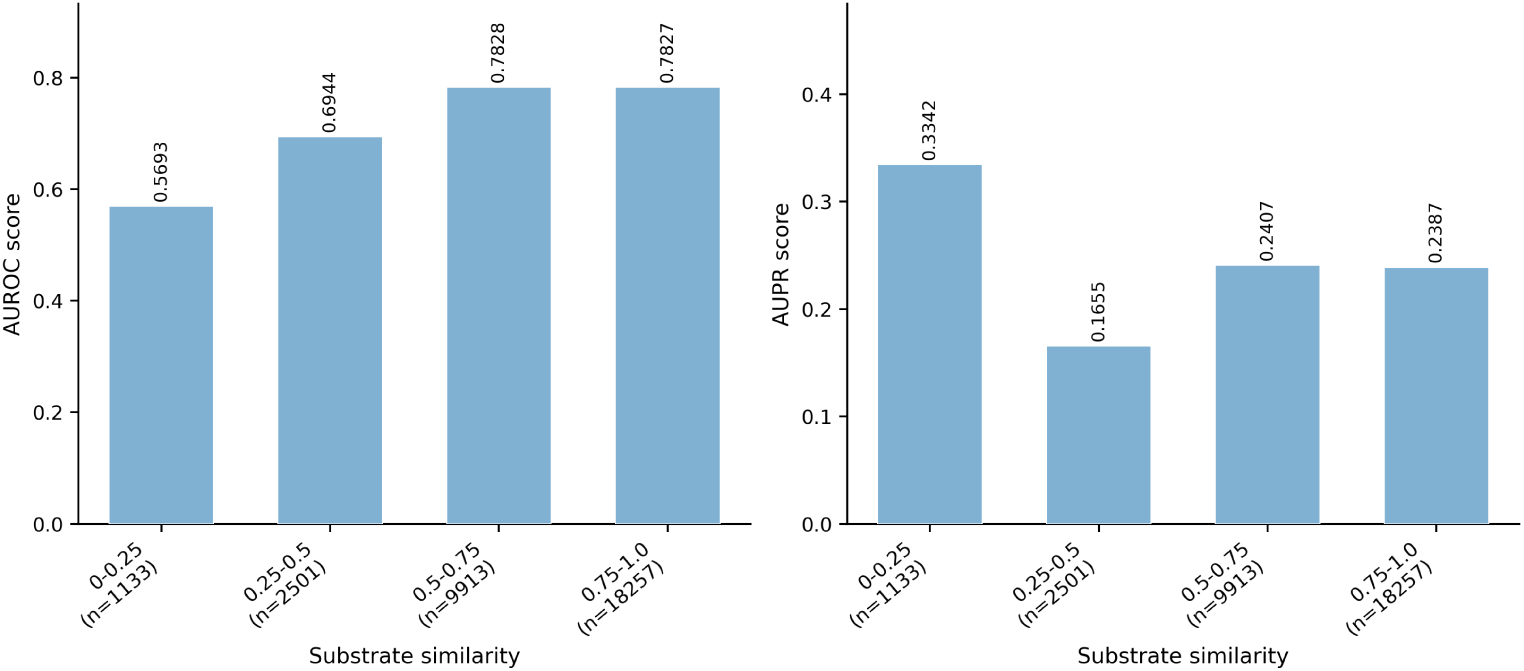
Boltz2ESI performance on the unknown substrate and enzyme split stratified by substrate novelty. Test-set enzyme–substrate pairs are binned by the maximum Morgan fingerprint Tanimoto similarity between the test substrate and any training-set substrate. AUROC (left) and AUPR (right) are shown for each bin, with sample sizes indicated in parentheses.

## Notes

### Competing Interest Statement

The authors have declared no competing interest.

